# Dengue virus infection changes *Aedes aegypti* oviposition olfactory preferences

**DOI:** 10.1101/375972

**Authors:** Julie Gaburro, Prasad N Paradkar, Melissa Klein, Asim Bhatti, Saeid Nahavandi, Jean-Bernard Duchemin

## Abstract

*Aedes aegypti* mosquitoes, main vectors for numerous flaviviruses, have olfactory preferences and are capable of olfactory learning especially when seeking their required environmental conditions to lay their eggs. In this study, we showed that semiochemical conditions during *Aedes aegypti* larval rearing affected future female choice for oviposition: water-reared mosquitoes preferred to lay eggs in water or p-cresol containers, while skatole reared mosquitoes preferred skatole sites. Using two independent behavioural assays, we showed that this skatole preference was lost in mosquitoes infected with dengue virus. Viral RNA was extracted from infected female mosquito heads, and an increase of virus load was detected from 3 to 10 days post infection, indicating replication in the insect head and possibly in the central nervous system. Expression of selected genes, potentially implied in olfactory learning processes, were also altered during dengue infection. Based on these results, we hypothesise that dengue virus infection alters gene expression in the mosquito’s head and is associated with a loss of olfactory preferences, possibly modifying oviposition site choice of female mosquitoes.

## Introduction

It is important for insects to identify and localize sites for food resources, mating and egg laying to successfully survive and reproduce. It is generally known that insects, including mosquitoes, are capable of information retention, acquired either during larval development, hatching or foraging, and that new information could influence their behaviour ^1^. In this study, we define the term of “olfactory learning” as change of olfactory preferences induced by exposure to odour.

In insect vectors, such as mosquitoes, learning from its own experience could influence the vector behaviour, such as the potential preferences in vector-host interactions^2,3^, which can be of great interest to epidemiologists. Mosquitoes are also capable of associative learning, which is the process of learning the association between two stimuli ^4^, in the context of feeding ^5^, or responding to odour cues ^6–10^. Also, learning in mosquito vectors could significantly influence virus transmission through host contact and survival rate ^9^. In nature, environments are constantly subjected to changes, especially chemical cues involved in potential food or oviposition site ^11^. In order to respond to such rapid change and improve its searching efficiency, a vector needs to adapt by learning to respond to odour cues ^6^. The mosquito experience to olfactory cues from larval, pupal or early adult vector environment could influence its behaviour.

After a blood meal from a host, female mosquitoes look for suitable oviposition sites to guarantee success of the offspring ^12^. Among different factors, chemical cues play a key role in determining location for mosquito to lay its eggs^13,14^. The chemical signature of an ideal oviposition site is not certain; thus, mosquitoes have to respond to rapid environmental changes to reduce risk of choosing an unsuitable site for their eggs by relying on previously experienced odour cues. Field studies have shown that mosquitoes have site-fidelity^15,16^, suggesting that early stage learning is important in resource location in adult mosquitoes. Processes of how and at which developmental stage mosquitoes acquire this information are still unknown and we do not address this question in our study. However, data showing olfactory site preferences after artificial odour induction during larval and pupal rearing in vectors^17,18^, suggest an important role of the mosquito olfactory learning in its oviposition choices ^19^.

It has been shown with *Culex quinquefasciatus* ^18^ and *Aedes aegypti* ^17^ species, that such olfactory learning of mosquitoes during their larvae and pupae development in different semiochemical environments can affect their oviposition choice and deterrence to those chemicals. Skatole, (3-methylindole) a naturally occurring compound in faeces, was prepared from fermented Bermuda grass and has strongly induced oviposition at concentrations ranging from 10^-5^ to 10^-3^ ppm in water for *Culex quinquefasciatus* ^20–23^. Skatole has also been successfully used in the field with oviposition bait trap sites in Tanzania for this mosquito species^24,25,^ however, higher concentration of the compound has shown a repulsive effect ^21^. *Culex quinquefasciatus* mosquitoes, initially repulsed by high concentrations of skatole, were shown to reverse their preference for oviposition sites when reared in skatole at the same concentration ^18^. The same behaviour was observed with *Aedes aegypti* conditioned with another repellent, Mozaway™ ^17^. In *Aedes* mosquitoes, skatole has been found to be an attractant at 100 ppm or higher concentrations ^26^, however, data is not available for lower concentrations. The semiochemical 4-methylphenol or p-cresol was found to be preferentially chosen by *Culex quinquefasciatus* females at specific concentrations^18,22^, but *Aedes aegypti* species shows contradictory data concerning oviposition attraction^27,28^. Semiochemical preferences vary with the mosquito species and concentration, (reviewed by Afify *et al*. ^29^), but also with experimental assay conditions, including rearing methods, or the presence of background odours ^30^.

*Aedes aegypti* is the main vector for dengue virus and is a major global public health problem ^31^. Some studies have reported behavioural changes due to arbovirus infection in mosquito vectors, such as during blood-feeding ^32–35^, host-seeking ^36^ and locomotion^37,38^. However, no study has reported a change in oviposition behaviour of the vector due to viral infection. Modification of vector preferences could act as potential drivers for virus range expansion, and hence its transmission, by increasing its vector distribution.

Studies on olfactory preferences in mosquito vectors are important for vector control strategies. Here we addressed the question of oviposition preferences of adult mosquitoes by semiochemicals rearing of larvae and possible alterations in olfactory preference after infection with dengue virus, here serotype 2 (DENV2). By using two independent behavioural assays, determining mosquito fitness, and dengue virus infection dynamics in their heads associated with selected gene expression, we increase our understanding of vector manipulation by arboviruses.

## Results

### Baseline oviposition choice

Initially, we tested the oviposition preference of *Aedes aegypti* females for skatole and p-cresol. We used a fixed concentration of 100 μg/l for all chemicals used in this study. As indicated on the experimental setup (Fig. 1A), female mosquitoes had 3 choices where to lay their eggs: either water, skatole or p-cresol. Our results showed that female mosquitoes had a significant repulsion towards skatole with less than 15% of eggs laid in this container (Fig. 1B). Females were indifferent to water or p-cresol with a nearly identical percentage of eggs laid in both water and p-cresol containers (respectively, 43 and 44 %).

**Figure 1:**
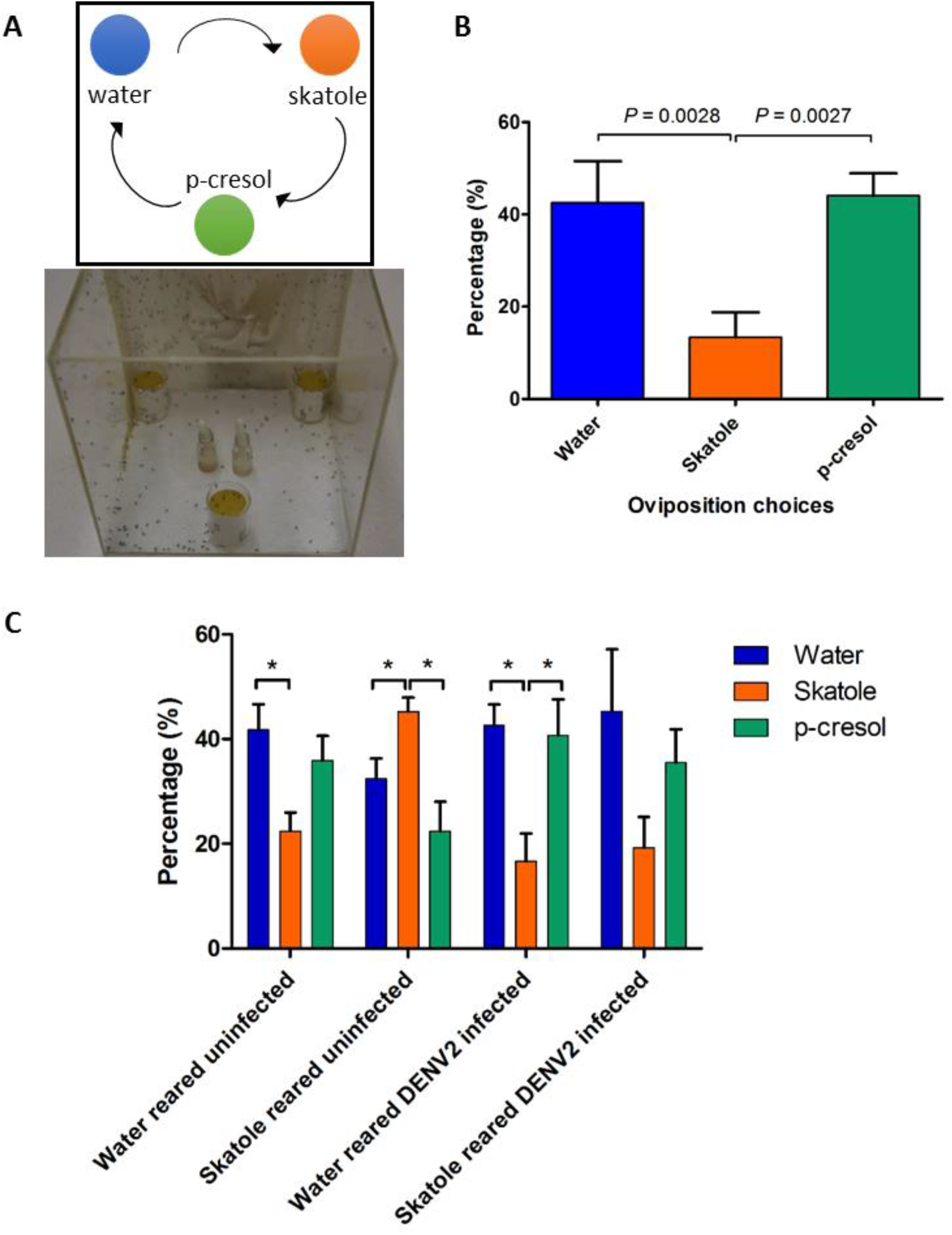
Oviposition choices after induction of olfactory preference in larvae and infectious blood-meal in adult *Aedes aegypti* females. (**A**) Illustration of the experimental setup for the preliminary assay on chemical preferences of the laboratory mosquito colony. Each of the three beakers contains either water, skatole or p-cresol chemicals (100 μg/l) with a sandpaper strip, for females to lay their eggs after a blood meal. The containers are equidistant and their position is daily moved clock-wise to avoid any location preference in the cage. (**B**) Results of the preliminary experiment with the *Aedes aegypti* colony, showing the percentage of eggs laid in each container after a blood-meal. (**C**) Bar plots showing water and skatole-reared female preferences in percentage of eggs in each container after an infectious blood-meal. Data shows the mean percentage (± sem) of the four experimental replicates and *P*-values for significant differences between the egg percentages with \**P* < 0.05; \*\**P* < 0.01; \*\*\**P* < 0.001.

### Oviposition choices post-skatole conditioning and DENV2 infection via a blood feeding

After blood-meal (BM), females reared in water showed similar results as the baseline experiment with a significant repulsion towards skatole container compared to water (Fig. 1C). On the other hand, females reared in skatole showed a significant attraction (Mann-Whitney test, *P* = 0.0286) to skatole container to lay their eggs compared to either p-cresol or water containers. These results show that females are influenced in their olfactory choice of the oviposition container by the conditions of their aquatic rearing medium. Blood-meal with dengue virus did not modify the oviposition choice of females reared in water; preferring to lay eggs either in water or p-cresol rather than skatole. However, in the case of skatole-reared females, the infection with DENV2 changed the oviposition choice. Instead of being attracted to skatole during oviposition, as with uninfected skatole reared group, infected females did not show any significant preference towards a specific container. Female mosquitoes were given a second BM and another choice of three containers to lay their eggs (Supplementary Figure S1). Uninfected and water-reared females showed a different pattern of olfactory choices compared to the first blood-meal with no significant preference between the three different containers, having lost the repulsion towards skatole. The attraction to skatole of the uninfected skatole-reared females was also weaker with an increase of eggs laid in p-cresol. Finally, water and skatole-reared females infected with DENV2, did not show any significant attraction or repulsion towards any container after a second BM, with no significantly different average number of eggs.

### Y-tube assays

After being released in the Y-tube olfactometer (Fig. 2A), mosquitoes displayed olfactory choices according to their respective rearing condition and infection status (Fig. 2B). Similar to the oviposition choice experiment, after the first blood-meal, uninfected females preferred the arm with the same odour stimulus where they were reared: females reared in water had a significant negative Preference Index (PI) compared to females reared in skatole (Mann Whitney test, *P* = 0.001), indicating preference for water over skatole. Independent of their rearing condition, dengue virus infected females preferred water for oviposition with a negative PI. As observed earlier, after the second BM, skatole conditioning disappeared in skatole-reared females with no preference between the skatole and water arms (Supplementary Figure S2). Furthermore, no preference was visible in skatole-reared and DENV2 infected females, confirming previous results with egg counting. From a binomial regression model analysis, we found that the most significant explanatory variables are, in decreasing order, the interaction infection status * reared condition (*P* < 0.0001), the interaction infection status * reared condition * days post infection (dpi) and reared condition alone (*P* <0.001). These results confirm that the changes in choice caused by DENV2 infection are correlated with larval rearing condition of the females.

**Figure 2.**
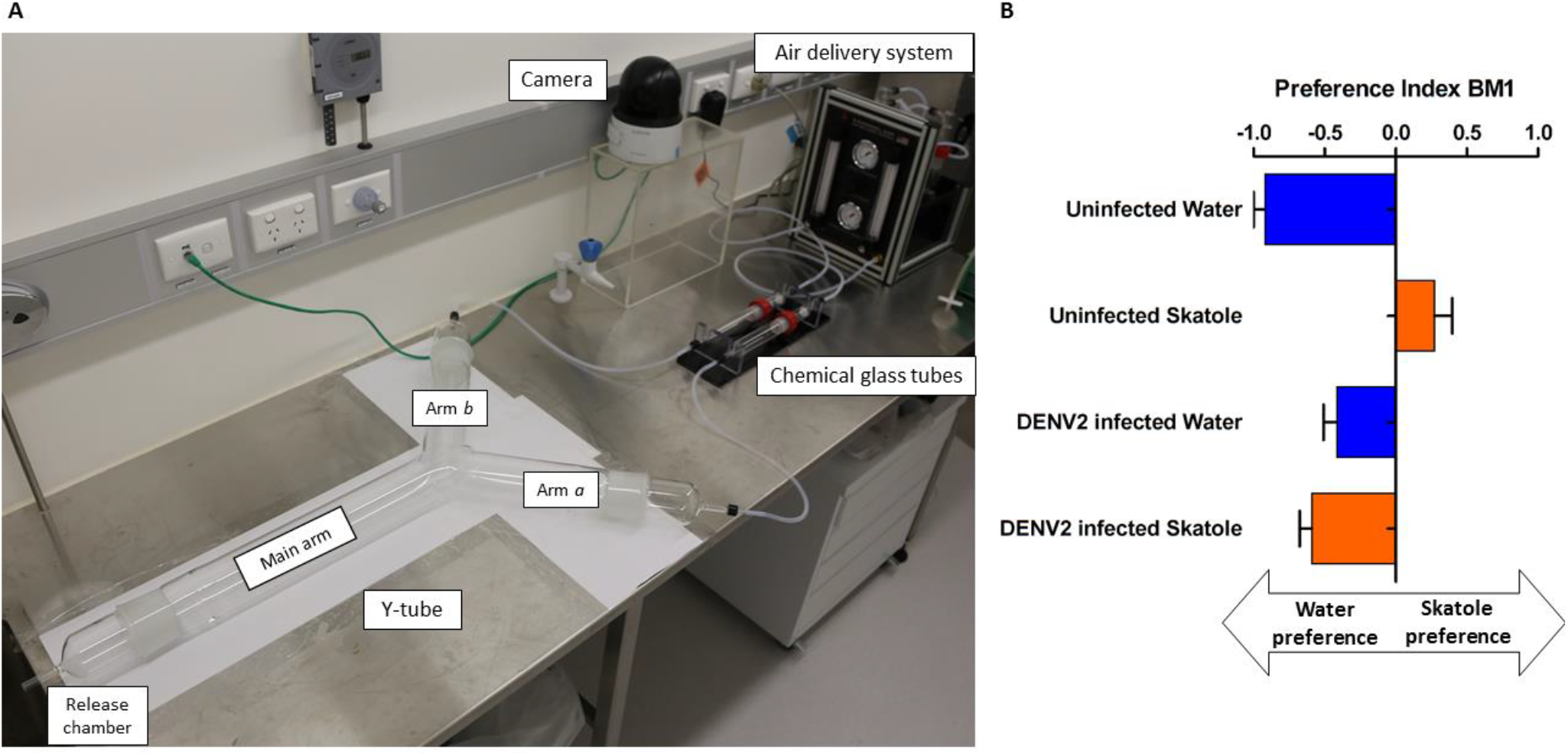
Dual olfactory choice between water and skatole during oviposition period of infected *Aedes aegypti* females. (**A**) Y-tube experimental setup: solvent (water) or skatole is place in one of the chemical glass tubes, leading to one of the arms of the Y-tube. The air delivery system allowed a controlled regular equal flow in each arm. Mosquitoes were released in the chamber and their location in the Y-tube recorded at the end of the assay. (**B**) Results of dual tests with different female groups after the first infectious blood-meal. The Preference Index (PI) indicates females that have chosen the arm with skatole (positive PI) or water (negative PI). Bar plots indicate means of PI (± sem).

### Infection dynamics and effect on female fitness

During the oviposition choice assays, the percentage of females blood-feeding during the first BM was recorded. There were no significant differences between the numbers of blood-fed females between the different conditions: rearing condition or DENV2 infection status (Table 1). During the oviposition choice experiments, the survival percentage of females (number of females alive after the second oviposition compared to the initial number post 1^st^ blood-meal) was not significantly different between the groups. The virus prevalence was confirmed in the DENV2 infected females used for oviposition choice assays. At 13 dpi, more than 80 percent of females were infected with DENV2 with no significant difference between water and skatole-reared females. Virus titration from the freshly blood-fed female abdomens, showed that each female took an average 1.22×10^6^ (± 3.66×10^5^) TCID_50_/ml of DENV2.

**Table 1:** Fitness parameter of female *Aedes aegypti* reared in water or skatole-reared and virus prevalence after DENV2 infection.

| Parameters | Group | Mean<br>(± sem) | Test ($N_{replicates}$)<br>*P* |
| --- | --- | --- | --- |
| Blood fed females during 1st BM (%) | Water uninfected | 28.25<br>($\pm 3.33$) | Friedman test<br>$F = 0.3$, ($N = 4$)<br>ns |
| | Skatole uninfected | 28.75<br>($\pm 4.8$) | |
| | Water DENV2-infected | 27.375<br>($\pm 7.52$) | |
| | Skatole DENV2-infected | 30.775<br>($\pm 5.8$) | |
| Female survival (%) | Water uninfected | 87.26<br>($\pm 4.56$) | Friedman test<br>$F = 2.45$, ($N = 4$)<br>ns |
| | Skatole uninfected | 89.73<br>($\pm 5.14$) | |
| | Water DENV2-infected | 84.17<br>($\pm 8.56$) | |
| | Skatole DENV2-infected | 79.39<br>($\pm 5.99$) | |
| Number of eggs laid post 1st BM (average per female) | Water uninfected | 82.38<br>($\pm 10.72$) | Friedman test<br>$F = 9.9$, ($N = 4$)<br>$P = 0.006$ |
| | Skatole uninfected | 88.52<br>($\pm 5.08$) | |
| | Water DENV2-infected | 68.73<br>($\pm 6.53$) | |
| | Skatole DENV2-infected | 60.47<br>($\pm 8.24$) | |
| Number of eggs laid post 2nd BM (average per female) | Water uninfected | 62.92<br>($\pm 13.25$) | Friedman test<br>$F = 1.8$, ($N = 4$)<br>ns |
| | Skatole uninfected | 67.93<br>($\pm 19.22$) | |
| | Water DENV2-infected | 52.41<br>($\pm 10.21$) | |
| | Skatole DENV2-infected | 57.85<br>($\pm 6.65$) | |
| Number of eggs laid for both BM (average per female) | Water uninfected | 74.04<br>($\pm 8.56$) | Friedman test<br>$F = 10.03$, ($N = 4$)<br>$P = 0.018$ |
| | Skatole uninfected | 79.70<br>($\pm 8.8$) | |
| | Water DENV2-infected | 61.74<br>($\pm 6.16$) | |
| | Skatole DENV2-infected | 59.35<br>($\pm 5.1$) | |
| Virus prevalence (%) | Water DENV2-infected | 93.31<br>($\pm 4.44$) | Mann Whitney test<br>$U = 3.0$, ($N = 4$)<br>ns |
| | Skatole DENV2-infected | 81.70<br>($\pm 0.86$) | |

Finally, the average number of eggs laid per female was estimated after counting the total number of eggs laid and dividing by the number of blood-fed females after each blood-meal. Although no significant difference was found between rearing condition groups, DENV2 infected females appeared to lay fewer eggs than uninfected mosquitoes after the first infectious blood meal (Table 2, *P* = 0.006).

**Table 2:**
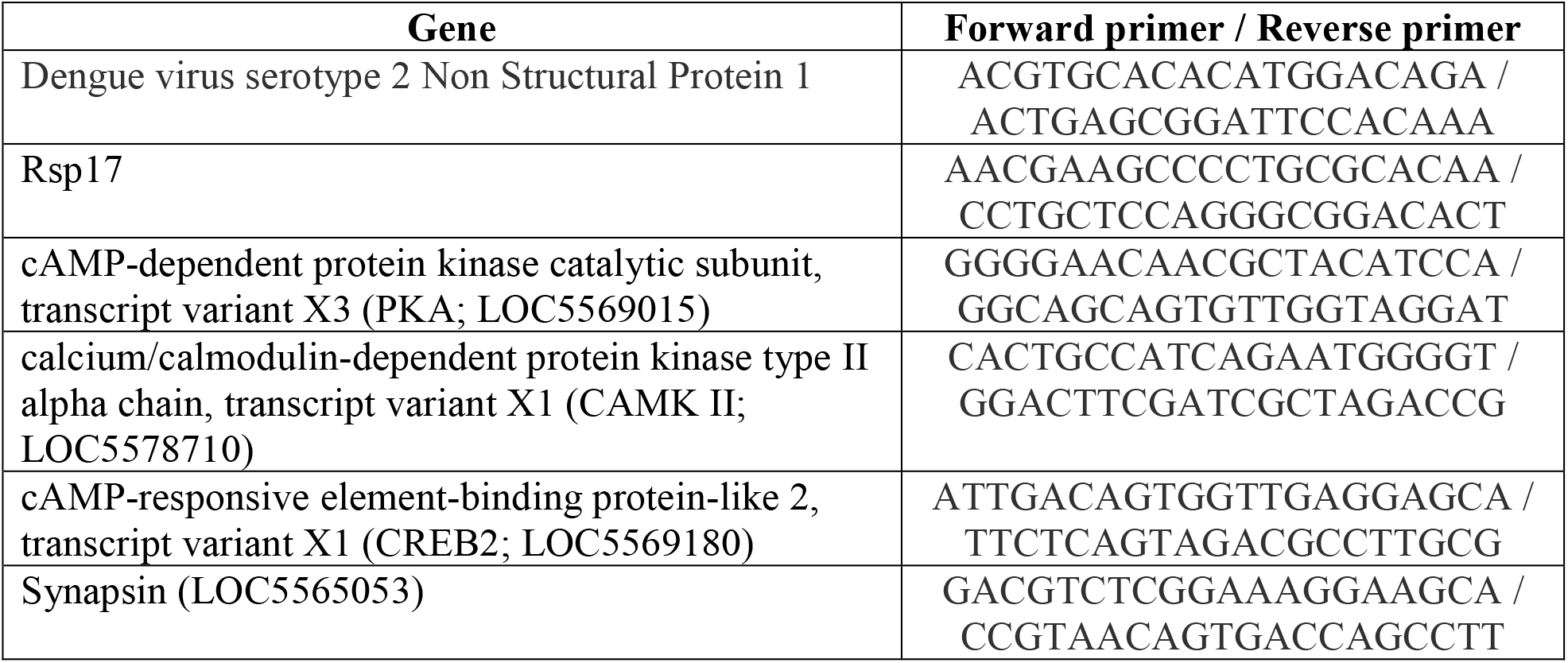
List of 5-3’ sequences of oligonucleotides primers for quantification of viral RNA and gene expression in *Aedes aegypti* heads.

### Virus replication and gene expression in mosquito heads

To determine when DENV2 reaches female mosquito’s head and potentially the central nervous system, RT-q-PCR tests were performed at different days post infection (dpi). Immediately after the blood feeding, the mosquitoes had detectable viral RNA from blood uptake in their heads, including the proboscis, with an average Ct of 33. Viral RNA in the mosquito head was then undetectable by day 3, before rising to detectable levels at 4 dpi. By 7 dpi, DENV2 RNA was detected in all tested heads, with an average Ct of 25. The amount of detectable RNA increased to an average Ct of 21 by 10 dpi representing a significant increase of virus in the mosquito heads (Fig. 3).

**Figure 3.**
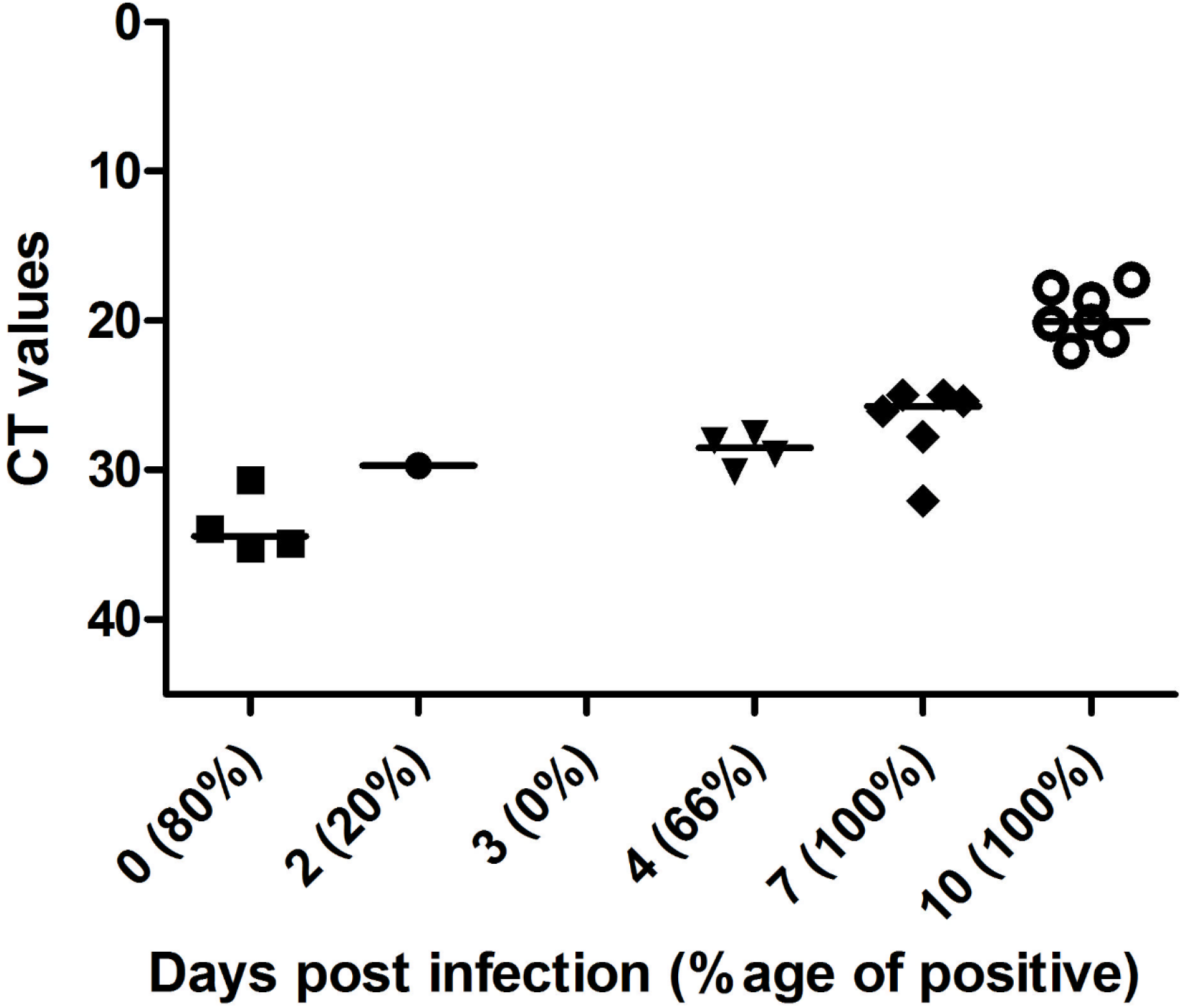
Dengue virus serotype-2 RNA in the heads of *Aedes aegypti* at different days post infection (dpi). Ct values for virus RNA detected via RT-q-PCR in the heads of infected females at different dpi. Note that the Y-axis is inverted and that negative values (> 45) are not shown. The prevalence of positive Ct values for DENV2 RNA is in brackets after the day post infection, on the X-axis.

To determine the effects of virus replication in the mosquito, we looked at the contemporary changes in gene expression in mosquito head during DENV2 infection. Two genes of the cAMP-responsive element-binding protein (CREB) pathway were targeted (Table 1), as they are broadly involved in synaptic plasticity and learning process in *Drosophila*, as well as *Aplysia* and rodent models ^39^. The cAMP-dependent protein kinase A (PKA) is activated by intracellular calcium increase following the activation of synaptic neurotransmitter receptor. Acting as the second messenger, cAMP phosphorylates the CREB, and is therefore an active transcription factor for CREB-dependent gene expression ^40^. Gene expression of the calcium/calmodulin-dependent protein kinase type II (CAMK II), involved in long-term potentiation, was explored (Table 2). This kinase is activated by calcium and calmodulin and is capable of auto-phosphorylation and controls neuronal growth in *Drosophila* ^41^. One mode of action of CAMK II is linked to phosphorylation of synapsin, a key-component of glutamate pre-synaptic vesicles, leading to their immediate availability as neurotransmitter pool ^42^. Expression profile for all four genes involved in learning or synaptic plasticity showed overexpression after infection compared with uninfected mosquitoes, becoming significant at 7 dpi (Fig. 4). However, a significant decrease was seen at 10 dpi for CAMK II (Fig. 4B) and Synapsin (Fig. 4D), compared with uninfected mosquitoes.

**Figure 4.**
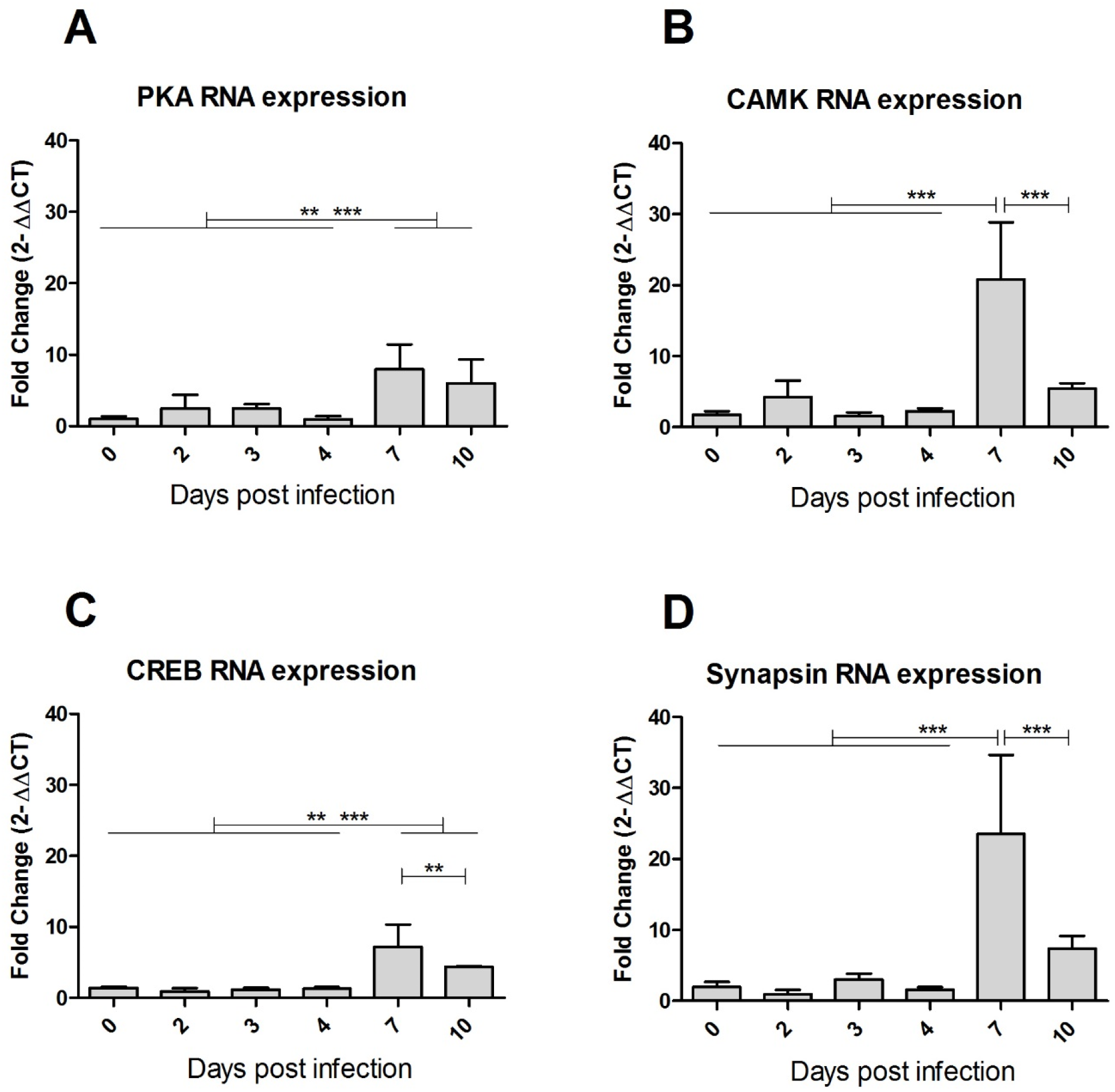
Gene expression in infected *Aedes aegypti* heads at different days post infection. Means with 95% confidence interval of RNA expression fold change detected via RT-q-PCR (2^ΔΔCt^ values, compared to Rsp17 housekeeping gene and to uninfected, for the corresponding day). Expression was analysed for the following genes: (**A**) cAMP-dependent protein kinase catalytic subunit (PKA), (**B**) calcium/calmodulin-dependent protein kinase type II alpha chain (CAMK), (**C**) cAMP-responsive element-binding protein-like 2 (CREB), and (**D**) Synapsin. *P*-values for significant differences between days with \**P* < 0.05; \*\**P* < 0.01; \*\*\**P* < 0.001.

## Discussion

Our behavioural assays demonstrated that female mosquitoes retain chemical information from the larval and pupal environment from which they emerge as adult. Understanding the developmental stage of when this happens and which cellular processes are involved in this behaviour, remain to be determined. This induced olfactory preference persists until the first oviposition, but fades after the second BM (Supplementary Figures S1 and S2), in the absence of later reinforcement ^19^. Despite being repulsive during baseline conditions, skatole at 100 μg/l could be attractive for gravid *Aedes aegypti* females if previously reared in water containing this chemical. These results verified that early experience of mosquitoes at aquatic stage could have an impact in the adult mosquito life^7,43^. As previously reported^15,16^ for *Anopheles* malaria vectors, the dissociation between oviposition site, which is often scarce, and remote food source implies needs for long or mid-range orientation and would necessitate “some form of learnt behaviour” ^15^.

In this study, the impact of DENV2 infection on its vector behaviour was tested on female mosquitoes reared in water or skatole. No significant change was observed after infection in water-reared females. Interestingly, after infection, skatole-reared mosquitoes did not show any preference for the chemicals (Fig. 1C) and preferred the water arm in the Y-tubes assays (Fig. 2B). However, after the second not-infectious BM in the oviposition choices experiment, no difference was observed between containers (Supplementary Figure S1), indicating that the skatole was not a deterrent at that stage. Interestingly, besides the observed behavioural changes, infected females also showed a significantly decrease number of eggs laid. This potential fitness cost has previously been reported in *Aedes aegypti* infected by DENV2 ^32^.

In parallel to the behavioural data, RT-q-PCR results showed that DENV2 RNA was present in the head of orally infected females immediately after an infected blood meal, probably through contamination of the buccal pieces and disappeared at 3 dpi. The prevalence of DENV2 positive mosquitoes then increased from 0 % at 3 dpi (Fig. 3) to 66 % of positive females at 4 dpi, and finally 100 % at 7 and 10 dpi. The increase in prevalence was associated with increase in viral RNA load in mosquito heads from 4 dpi to 10 dpi (Fig. 3), probably due to virus replication. Previously, dengue virus antigen detection in orally infected *Aedes aegypti* mosquitoes showed that the nervous tissues are among the first to be infected, as early as 5 dpi ^44^. In this study, the authors highlighted that DENV2 (Jamaica strain) infection was absent in the *Aedes aegypti* (Mexico origin) muscles, unlike previously shown *Culex pipiens* infected with West Nile virus ^45^. Interestingly, these authors also stated that heads and salivary glands were the only tissues where viral antigens continued to accumulate until 21 dpi, while other infected tissues showed a decrease in the infection rate. Finally, a recent study showed that the same DENV2 strain is able to replicate in dissected *Aedes aegypti* (Mexico origin) primary neuron culture, with the presence of viral antigens ^38^. Oviposition behaviour, as well as the related odour choice modification by infection, occurred between 3 to 5 days post BM ^46^, with increase of virus load and prevalence in female heads. Therefore, it can be hypothesised that DENV2 may interfere with the olfactory signal processing, either involving the sensory organs (antennas, palps) or potentially involving neurons from the central nervous system, as postulated for *Culex pipiens* infected with West Nile virus, which displayed behavioural changes post infection ^36^.

At the molecular level, many genes have been implicated in olfactory learning using the *Drosophila* model (for a full list of genes, see review ^47^). Some of these genes encode components in the cAMP signalling pathway, such as the cAMP-dependent protein kinase (PKA) and the transcription factor cAMP-response element binding protein (CREB). We investigated the local effect of virus replication in naturally infected mosquito brain and our results showed overexpression of PKA and CREB genes in tested mosquito heads, peaking at 7 dpi. *Aplysia, Drosophila*, and rodent models have shown that CREB-dependent transcription was required for cellular mechanisms underlying long-lasting learning ^48^. In addition, these molecular learning mechanisms revealed to have a degree of conservation across widely evolved species ^39^. In insects, the PKA/CREB pathway also participates in the transcriptional enhancement by the steroid hormone, ecdysone and its 20-Hydroxyecdysone metabolite ^49^, which is overexpressed in mosquitoes temporarily, peaking 24 hours after blood-meal. These hormones are involved in long term courtship neuro-processes in *Drosophila* ^50^. In mosquito cell culture, transcriptome analysis has shown over-expression of CREB after WNV infection ^51^. CREB can be phosphorylated by PKA, CAMK and microtubule associated protein kinase (MAPK) ^52^. However, given the wide range of processes involving cAMP and CREB pathways, notably during the development, we cannot rule out the probability of gene expression modification due to other cell types (haemocytes) or organs (neuro-endocrinal glands) also present in the head, not belonging to the nervous system. As previously mentioned, muscle tissues could reasonably be considered not affected by the infection and it is noteworthy, that the nervous system is a major component of the mosquito head, in terms of cell quantity. Our mRNA expression data, although limited, showed elevated mRNA levels for CAMK in mosquito heads after DENV2 infection. CAMK II and its auto-phosphorylation capacity has been found to be pivotal in *Drosophila* learning processes ^41^. This kinase has also been shown to phosphorylate synapsin I ^53^, which is pivotal in olfactory learning in *Drosophila* ^54^. Although the calmodulin pathway is also involved in development processes, particularly for the nervous and visual systems, and the immune system of invertebrates; synapsin, on the other hand, is more specific to neuronal cells. Our data shows that dengue virus infection in mosquito heads led to overexpression of genes involved in memory, some of which are also implicated in immune defence in mammals ^55^. This overexpression coincided with the presence of DENV2 RNA in the heads of infected mosquitoes. Behavioural changes observed at 4-5 dpi, occurred slightly before significant vector gene expression change. We cannot rule out the possibility that the viral infection impacts on tissues other than the central nervous system, like buccal pieces, neuro-endocrinal glands, antenna and eyes. These could be hypothetically involved in the observed gene expression variations and behavioural changes, and further research is needed to identify appropriate tissue involved. The dynamics of behaviour, viral prevalence and molecular changes in the mosquito were associated in time in the mosquito head, leading to the hypothesis of virus-induced molecular changes in the pathways potentially involved in mosquito olfactory learning.

Although no previous experiments have reported modifications in oviposition choices in DENV2 infected mosquito females, other behavioural changes have been described. *Aedes* mosquitoes after artificial intrathoracic infection showed an increased total level of activity 2 to 6 dpi, with similar daily activity patterns between DENV2 infected and uninfected mosquitoes ^37^. Another study showed that infected Aedes aegypti females infected with DENV2 had an increased activity and space occupation in laboratory conditions ^38^. Behavioural changes post infection underlie a potential way for the virus to manipulate the vector. Thus, viral transmission range could possibly increase, by widening space exploration of infected females during oviposition behaviour. Although we cannot exclude detrimental effect of infection on the vector’s fitness with lower egg production and potential altered choice for larval rearing medium, these modifications in the infected mosquito could be advantageous for the virus by broadening the spatial range of vectors and colonizing new ecological niches.

## Conclusions

In this study, we examined “olfactory learning” behaviour of the mosquito, *Aedes aegypti*, defined as a change of olfactory preferences induced by skatole exposure. Our results showed a change in oviposition preference of *Aedes aegypti* caused by DENV2 infection. Based on association of gene expression data with the behavioural changes and previously published data on these genes, we believe that DENV2 infection leads to mosquito learning behavioural changes via changes in these gene expression. However, we cannot rule out the possibility that these genes are involved in a different type of learning, such as habituation, or associative learning. These behavioural changes could have an impact on viral transmission, as they could be either advantageous for virus dispersion via infected vectors, or detrimental for the vector’s fitness. These results showed that, far from being neutral and innocuous to the vector, the virus infection triggered complex changes in the mosquito behaviour and its potential fitness. It is also important to note that only genes involved in learning processes were assessed, and that genes involved in chemosensory, neuronal circuit development or other neuronal circuit pathways could also be involved. The findings presented here are important for dengue control strategies in order to find optimal strategies for adult/larvae trapping and insecticides treatment ^56^.

## Material and Methods

### Mosquito colonies and induction of olfactory preferences in larvae

All experiments were conducted in a PC3 insectary with *Aedes aegypti* colony, collected in December 2014 in Brisbane, Queensland. Mosquitoes were maintained at 28 °C with 70% humidity and at a photoperiod of 10:10 light:day (L:D) cycle with 2 hours transition of dim light in between. Eggs on paper sand strips were collected from the colony cages, dried and stored at room temperature.

About 400 eggs were allowed to hatch in either the control condition (water + solvent) or induced olfactory preference (water + skatole), in the same environmental conditions. After hatching, larvae were reared for about two weeks with equal repartition of fish food. After the apparition of the first pupae in the larval container, pupae and larvae were placed in an emergence cage with their respective rearing media. After emergence, adults were regularly collected and transferred in separate plexiglass cages (30 × 30 × 30 cm). Males and females were kept together in the cage for 4-5 days for mating. Mosquitoes had access to sugar (10% solution in water) *ad libitum*. One day before the BM, females were counted and equally distributed in four plexiglass cages corresponding to four experimental conditions: “Water reared uninfected”, “Skatole reared uninfected”, “Water reared DENV2 infected” and “Skatole reared DENV2 infected”. Experiments were performed in four replicates.

### Dengue virus infection via blood meal

Dengue virus serotype 2 ET300 (DENV2) used for our experiments was isolated in Australia from a soldier returning from East Timor and was obtained from Queensland Health. The virus was passaged seven times in *Aedes albopictus* C6-36 cell line. To prepare the blood solution, 1 ml of fresh chicken blood was centrifuged and plasma replaced by either L15 media for uninfected blood (control), or with 500 μl of DENV2 virus solution at a titre of 10^7^ TCID50/ml.

Female mosquitoes were fed on chicken blood using an artificial membrane feeding system (Hemotek system) utilising chicken skin. After the first infectious BM, only blood-fed females were selected for further studies. To improve the infection rate of females, the temperature was raised from 28 to 30 °C for two days following the first BM, as previously shown ^57^. Females were allowed 6 days to lay their eggs on sandpaper strips in clean glass beakers filled with the different semiochemical conditions. A second non-infectious BM was given at 7 days post infection (dpi).

### Semiochemical preparation

Semiochemicals, p-cresol and skatole (Sigma-Aldrich), were each dissolved in 70% ethanol to create a stock solution (1 mg/ml_ethanol_) and then diluted at 100 μg/l in 400 ml water for larvae and pupae conditioning. The high concentration allows for constant exposure of the aquatic stages of the mosquitoes during the rearing time.

### Behavioural assays

#### Baseline experiments

To check oviposition preferences before infection, preliminary bioassays were conducted with the *Aedes aegypti* mosquitoes using three olfactory oviposition choices. In short, after a routine blood meal, the colony cage was offered three different choices where the females can lay their eggs for 5 days *post* blood-feeding. Eggs were collected and counted for each condition ^28^.

#### Olfactory oviposition choices

For each experimental assay, each cage contained between 20-60 blood-fed *Aedes aegypti* females. After the blood meal, at 2 dpi to 6 dpi post first BM and from 9 to 13 dpi after second BM, females were given a choice for three olfactory oviposition containers to lay their eggs: water, p-cresol and skatole at the same concentration as larval rearing. The containers were placed at equidistance length from each other in the cages (2 in the corner and 1 against the middle of cage face), and rotated daily clockwise to avoid site preference. Eggs laid on the sandpaper strips within containers were collected at 6 dpi and 13 dpi and allowed to dry out at room temperature. Images from the sand paper strips were taken using a Leica microscope (DFC425) with 8.0× magnification for egg counting with Icount software ^28^. Mosquitoes from DENV2 infected groups were sacrificed after the experiment to check for proportion of infected mosquitoes. For each of the four replicates, the percentage of female blood-fed during the first BM and survival rates were recorded.

#### Y-tube assays

To test the dual-olfactory preference between water and skatole after olfactory preference induction in larvae and/or DENV2 infection, we used a Y-tube olfactometer (see setup Fig 2A) at different days post BMs. Data analysis did not reveal any bias towards the left or right side of the olfactometer (Mann-Whitney U-test, *P* = 0.94). Females were infected the same way as the oviposition choice experiment, however no container was provided to allow laying of eggs before the commencement of the experiments. Five to 12 females were placed in the release chamber, at the top of the long leg of the ‘Y’, and allowed to fly for 10-15 minutes. During data sampling, females were marked as “inactive” if they did not leave the release chamber, “active” if still in the main arm or “arm a” or “arm b” if they had chosen any of those arms. Females were placed back in their corresponding group cage and another set of females was randomly chosen for the next assay. In total, 3 repetitions per day post infection were performed for each group at 4 and 5 dpi for the first BM and 11 and 12 dpi for the second BM. We used a preference index (PI) to visualize the dual-choice as described by Vinauger *et al*. (2014) ^9^:

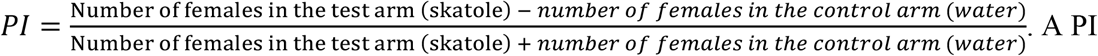

of +1 indicated that all females chose the arm_skatole_, a PI of 0 indicated that half of females chose the arm_skatole_ and half the arm_water_, and a PI of –1 indicated that all females chose the armwater of the olfactometer.

### Virus prevalence in infected groups – TCID50

At the end of the behavioural assay all surviving females from infected groups were collected in individual tubes, stored at −80 °C and tested in end point titrations (TCID50) using Vero (ATCC CCL81 African green Monkey Kidney) monolayer cell cultures assays to determine the overall percentage of infected females in each group condition. Abdomens from 10 freshly blood-fed females were used to measure the starting virus load taken during infectious BM. At the end of the experiment, dried bodies/abdomens were cleaned with 70% ethanol for 5 minutes. The rinse solution was removed and 150 μl of L15 media with 3-5 silica beads were added to each tube. Bodies/abdomens were homogenized for 5 minutes with a BeadBeater. The samples were centrifuged for 3 minutes at 11000 RPM. The supernatants were diluted 10-fold and tested in triplicate in a TCID50 procedure using Vero cells and incubated at 37°C with 5 % CO2. Cytopathic effect at 3 and 5 dpi was measured to determine TCID50 using Spearmann-Karber calculation method ^58^.

### Quantitative RT-PCR for virus detection and gene expression in mosquito heads

After an infectious BM, 5 to 7 female heads were sampled at 0 dpi (a few minutes after the BM), 2, 3, 4, 7 and 10 dpi and stored in RLT-buffer with 5-10 silica beads at −80 °C before testing. Total RNA was extracted from the samples using RNeasy Plus Mini Kit (Qiagen Sciences, Maryland, MA) and cDNA was prepared using random hexamers and Superscript-III reverse transcriptase (Thermo Fisher Scientific Inc. Australia) as per manufacturer’s protocol. Real-time PCR assay was performed using the SYBR^®^ Premix Ex Taq™ II (Takara-Bio Inc, China) on a QuantStudio™ 6 Flex Real Time PCR System (Applied Biosystems). The list of primers used for DENV2 RNA quantification and gene expression are listed in Table 2. Cycling was as follows: 95°C for 30 seconds, followed by 45 cycles of 95°C for 5 seconds, 55°C for 40 seconds, followed by melt-curve analysis. The cycle threshold (Ct) values, in duplicate for each sample, were collected at each time point for each condition. For RNA expression, the 2^ΔΔCt^ values were calculated at each time point for each gene as the fold-increase over uninfected control and normalized to Rsp17 housekeeping gene expression at the same time point. Ct values were measured at 0, 2, 3, 4, 7 and 10 dpi for 3 to 6 mosquitoes at each time and compared to control values of uninfected mosquitoes at the same time.

### Statistical analysis

Graphs were plotted and statistical analysis performed using GraphPad Prism 5 software. All tests were performed with a two-tailed analysis and graphs show the average with standard error of the mean (sem) (Fig. 1 and Fig. 2) or standard deviation (sd) (Fig. 4) as error bars. For Y-tube assays, a generalised linear mixed model (GLMM) with a binomial distribution and logit link function was additionally used to test for the effect of the different parameters (rearing condition, infection status and dpi) on the PI ^59^. The interaction between each parameter was included in the model as fixed effects: glm(choice ~ reared conditions * infected status * dpi, family=binomial(link=‘logit’). For this analysis, the software R version 3.4.1 ^60^ was used and the function “Analysis of Deviance for Generalized Linear Model Fits” ^61^ to validate our model.

## Acknowledgements

We thank Peter Mee for his help in ordering and organizing the shipping of the Y-tube maze. We are also thankful to Queensland Health for providing the dengue virus strain.

## Data availability

Data are available upon reasonable request to the corresponding author.

## Author contributions

Experimental design: J. G.; Behavioural assays: J. G. and M. K.; Virology: J.G., P. N P. and J-B. D. Analysis: J.G. and J-B. D.; Writing - original draft: J. G.; Writing - review & editing: J.G., J-B. D., P. N P., A. B., S. N.; Supervision: J-B. D. and A. B.

